# Artificial Intelligence Enables the Label-Free Identification of Chronic Myeloid Leukemia Cells with Mitochondrial Morphological Alterations

**DOI:** 10.1101/2023.07.26.550632

**Authors:** Kohjin Suzuki, Naoki Watanabe, Satoru Torii, Satoko Arakawa, Kiyosumi Ochi, Shun Tsuchiya, Kazuhiro Yamada, Yoko Kawamura, Sadao Ota, Norio Komatsu, Shigeomi Shimizu, Miki Ando, Tomoiku Takaku

## Abstract

Long-term tyrosine kinase inhibitor (TKI) treatment for patients with chronic myeloid leukemia (CML) causes various adverse events. Achieving a deep molecular response (DMR) is necessary for discontinuing TKIs and attaining treatment-free remission. Thus, early diagnosis is crucial as a lower DMR achievement rate has been reported in high-risk patients. Therefore, we attempted to identify CML cells using a novel technology that combines artificial intelligence (AI) with flow cytometry and investigated the basis for AI-mediated identification. Our findings indicate that *BCR-ABL1*-transduced cells and leukocytes from patients with CML showed significantly fragmented mitochondria and decreased mitochondrial membrane potential. Additionally, *BCR-ABL1* enhanced the phosphorylation of Drp1 via the mitogen-activated protein kinase pathway, inducing mitochondrial fragmentation. Finally, the AI identified cell line models and patient leukocytes that showed mitochondrial morphological changes. Our study suggested that this AI-based technology enables the highly sensitive detection of *BCR-ABL1*-positive cells and early diagnosis of CML.

## Introduction

Chronic myeloid leukemia (CML) is a myeloproliferative neoplasm caused by the translocation of chromosomes 9 and 22, resulting in the Philadelphia chromosome^1, 2^, which leads to the formation of a *BCR-ABL1* fusion gene. CML has an annual age-standardized incidence rate of 0.75/100,000 cases globally^3, 4^. CML is characterized by monoclonal neoplastic proliferation of hematopoietic stem cells at each stage of blood cell maturation^5^, mainly in granulocytes. CML progresses through chronic, accelerated, and blastic phases; tyrosine kinase inhibitors (TKIs) are the primary treatment strategy for this condition^6^. Despite significant improvements in the prognosis of CML due to TKIs, only 40% of patients achieve treatment-free remission (TFR) after treatment cessation^7^ because of TKI-resistant *BCR-ABL* mutations^8, 9^ and remaining CML stem cells^10^. Long-term use of TKIs is associated with various adverse events such as cardiovascular complications, edema, rashes, fatigue, and pleural effusions^11^. Guidelines for discontinuing TKI treatment include the chronic phase, a low prognostic score, more than 5 years of TKI treatment, and maintained deep molecular response (DMR) for several years before discontinuation^12, 13^. Early treatment responses are considered to result in deeper molecular responses and higher DMR achievement rates in patients to increase the actual number of patients in TFR. Hence, the factors considered for current prognostic scoring tools (including Sokal^14^ and ELTS^15^ score), spleen size, peripheral blast count, and platelet count, increase with delayed diagnosis. Therefore, we hypothesized that early diagnosis and treatment initiations are important to improve the TFR rate^16^.

Flow cytometry is a useful tool for detecting various cells, including leukemia cells^17^; however, there are still technical limitations, such as the requirement to stain surface antigens and the inability to detect small amounts of leukemia cells in the absence of leukemia cell-specific markers. In particular, early detection of peripheral CML cells is challenging because few CML cell-specific surface antigens and no morphological changes have been reported previously. Thus, we proposed using a novel technology that combines artificial intelligence (AI) with flow cytometry to detect CML cells with high accuracy.

Ghost cytometry (GC) is a recently developed machine vision-based technology based on flow cytometry and AI^18^, which utilizes a micro-patterned illumination of light to obtain temporal waveform signals that reflect minute structural changes in cells. It was shown that machine learning-based analysis of the signals can identify various cell types without staining. GC has already identified peripheral blood leukocytes with comparable accuracy to surface marker-based classification.^19^; however, the mechanism underlying the identification of unstained cells using GC has not been completely elucidated. Moreover, although GC has shown promise in identifying various types of cells, its clinical utility remains unclear.

Recently, AI has been used in various fields of medicine; however, many challenges need to be addressed, especially the problem of not knowing the basis for AI-associated decisions^20^. Therefore, this study aimed to utilize this novel AI-based technology to identify CML cells in a label-free manner and investigate the morphological changes to elucidate the basis for AI-based identification.

## Results

### AI-based identification of BCR-ABL1-transduced cells

As an initial step to determine whether CML cells could be identified using AI, cultured *BCR-ABL1*- transduced cells were evaluated. Specifically, the *BCR-ABL1* gene was transduced into the erythropoietin (EPO) growth-dependent UT-7/EPO cell line derived from the acute myeloid leukemia (AML) M7, and the performance of AI-based identification was assessed against gene-untransduced UT-7/EPO cells. As a result, the two cell types were identifiable with an area under the curve (AUC) of 0.87 (Fig. 1A).

**Fig. 1.**
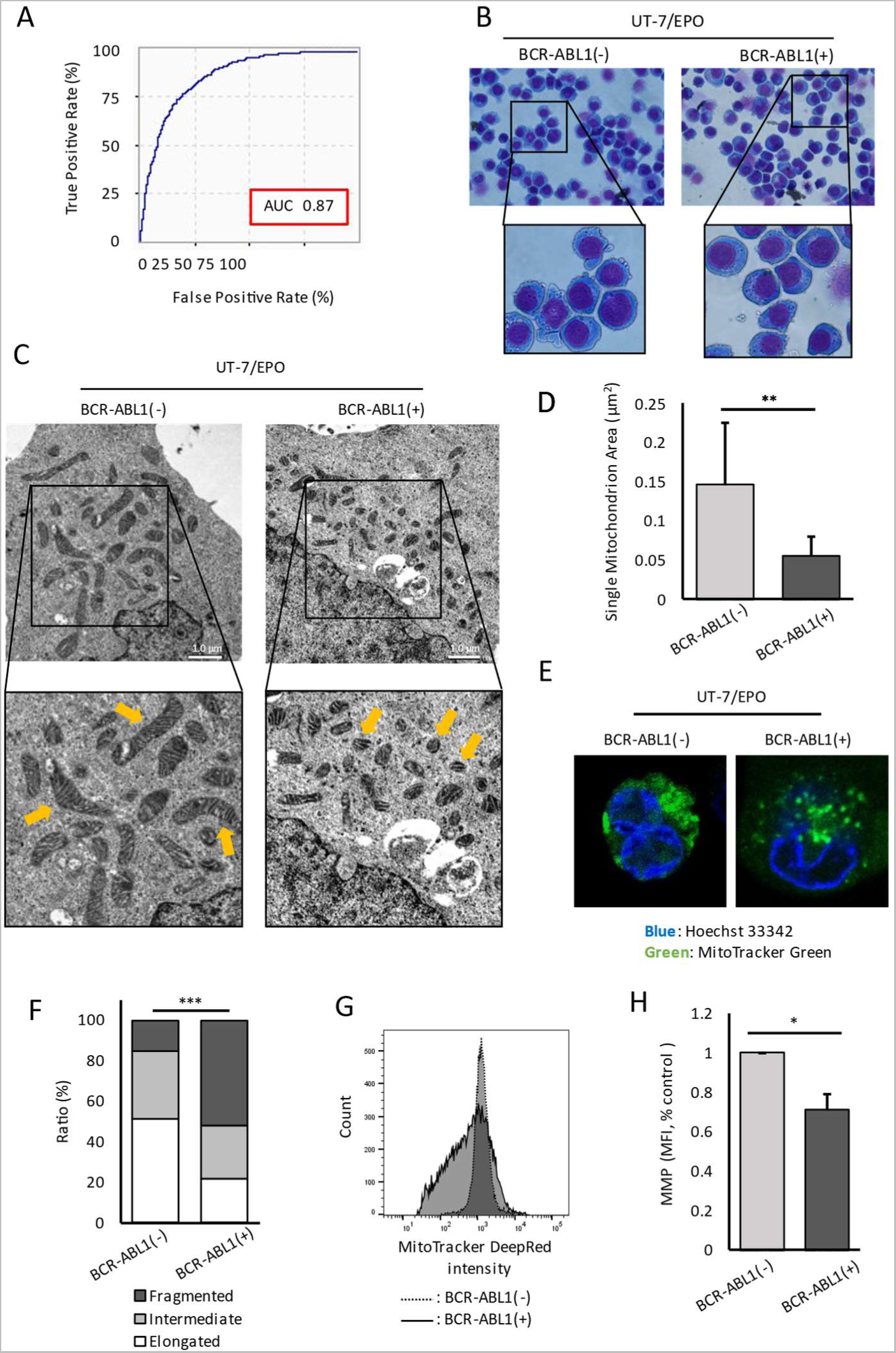
Examination of *BCR-ABL1*-transduced UT-7/EPO cells. A. An ROC curve and an AUC value for the AI-based identification of UT-7/EPO cells with or without *BCR-ABL1*. B. Typical Wright–Giemsa staining images of UT-7/EPO with or without *BCR-ABL1*. C. Typical TEM images of UT-7/EPO with or without *BCR-ABL1*. The white scale bar represents 1.0 µm. D. Quantitative results of a single mitochondrion area from the TEM images. Values are presented as the mean ± SD, **p < 0.01 by Mann–Whitney U test from three different experiments. E. Representative confocal laser scanning microscopy images of UT-7/EPO with or without *BCR-ABL1*. Mitochondria are shown in green and nuclei are shown in blue. F. The proportions of cells with different mitochondrial morphologies in UT-7/EPO with or without *BCR-ABL1*. Among the cells imaged using confocal laser scanning microscopy, the sizes of mitochondria are classified as Elongated, Intermediate, or Fragmented, and their respective ratios are presented. ***p < 0.001 by Fisher’s exact test. G. Typical flow cytometry histogram of UT-7/EPO with or without BCR-ABL1 stained by MitoTracker DeepRed. H. MMP measurement from UT-7/EPO cells. The average MFI of MitoTracker Deep Red staining from three different experiments is compared. Values are shown relative to those for *BCR-ABL1*(-). Values are presented as the mean ± SD, *p < 0.05 by Mann–Whitney U test. ROC, receiver operating characteristic curve; AUC, area under the curve; AI, artificial intelligence; *BCR-ABL1*(-), *BCR-ABL1*-untransduced; *BCR-ABL1*(+), *BCR-ABL1*-transduced; TEM, transmission electron microscopy; MMP, mitochondrial membrane potential; MFI, median fluorescent intensity.

Since our AI technology is reported to identify cells by their morphological differences^18^, the high AUC value obtained for the identification of *BCR-ABL1*-transduced UT-7/EPO cells indicates that the *BCR-ABL1* gene caused morphological changes in the cells; however, observation using Wright–Giemsa staining did not reveal any morphological changes (Fig. 1B). This suggests that the introduction of *BCR-ABL1* may result in intracellular structural changes that are not easily detectable using conventional microscopy-based examination. We then investigated the two cell types using transmission electron microscopy (TEM) and found that mitochondria were more fragmented in *BCR-ABL1*-transduced cells than in untransduced cells (Fig. 1C). Quantification of the mitochondrial area using the TEM image analysis showed a significant reduction in the area per mitochondrion in the *BCR-ABL1*-transduced cells compared with that in the untransduced cells (p < 0.01, Fig. 1D). In addition, observing the mitochondria in these cells stained with fluorescent MitoTracker Green revealed that *BCR-ABL1*-transduced cells exhibited mitochondrial fragmentation (Fig. 1E); furthermore, a significantly higher proportion of fragmented mitochondria was observed in *BCR-ABL1*-transduced cells compared to untransduced cells (p < 0.001, Fig. 1F).

To further analyze the change in mitochondrial function, we measured the mitochondrial membrane potential (MMP) of the two cell types by MitoTracker DeepRed and found a significant decrease in the MMP in *BCR-ABL1*-transduced cells compared with untransduced cells (p < 0.05, Fig. 1G, H). These findings suggest that *BCR-ABL1* induces mitochondrial fragmentation in UT-7/EPO cells, reducing MMP.

### Drp1 is implicated in the process of mitochondrial fragmentation in BCR-ABL1-transduced cells

After observing mitochondrial fragmentation in *BCR-ABL1*-transduced cells, we analyzed the changes in the expression of proteins responsible for regulating mitochondrial morphology. Our findings indicated that the expression levels of the mitochondrial fission factor dynamin-related protein (Drp)1, phosphorylation of Drp1 at serine 637 (Ser637), fusion factor mitofusin (Mfn)1/2, and optic atrophy (OPA)1 were not altered in *BCR-ABL1*-transduced cells compared with those in untransduced cells, except for the phosphorylation of Drp1 at serine 616 (Ser616), which increased in *BCR-ABL1*-transduced UT-7/EPO cells (Fig. 2A). Because p-Drp1(Ser616) is known to directly stimulate mitochondrial fission, its increase by *BCR-ABL1* is consistent with the morphological finding of more fission mitochondria.

**Fig. 2.**
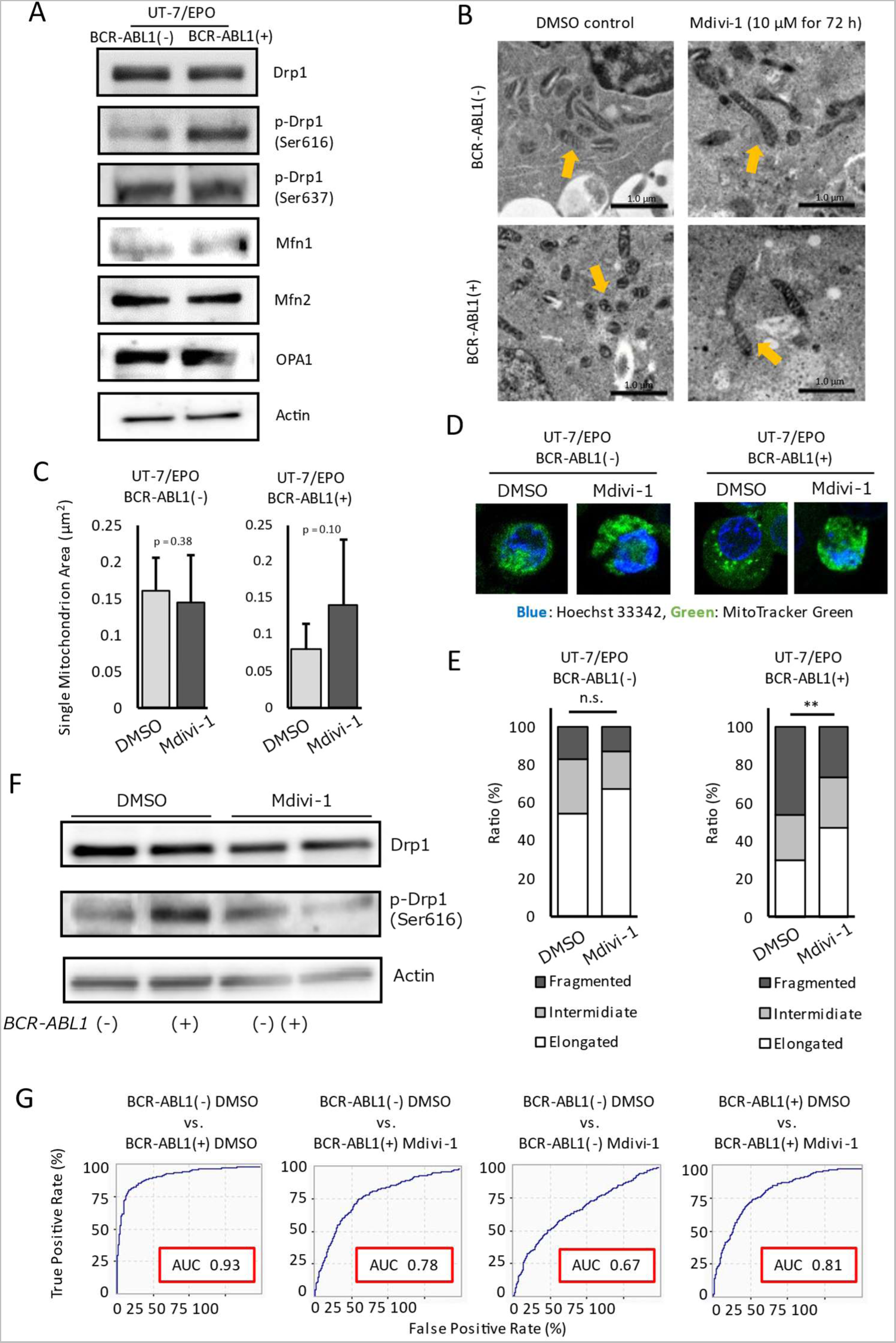
Drp1 is involved in mitochondrial fission by *BCR-ABL1.* A. Protein expression of UT-7/EPO cells with or without *BCR-ABL1*. B. Typical TEM images of UT-7/EPO with or without Mdivi-1 treatment (10 µM for 72 h). The black scale bars represent 1.0 µm, and the yellow arrows show the mitochondria. C. Quantitative results of single mitochondrion area of UT-7/EPO cells with or without *BCR-ABL1*, treated with Mdivi-1 from TEM images. P values were examined using the Mann–Whitney U test. D. Representative confocal laser scanning microscopy images of UT-7/EPO with or without *BCR-ABL1* treated with Mdivi-1 (10 µM for 72 h) or DMSO (control). Mitochondria are shown in green, and nuclei are shown in blue. E. The proportions of cells with different mitochondrial morphologies in UT-7/EPO with or without *BCR-ABL1* treated with Mdivi-1 (10 µM for 72 h) or DMSO (control). Among the cells imaged using confocal laser scanning microscopy, the sizes of mitochondria were classified as Elongated, Intermediate, or Fragmented, and their respective ratios are presented. **p < 0.01 by Fisher’s exact test. F. Protein expression of UT-7/EPO cells with or without Mdivi-1 treatment (10 µM for 72 h). G. ROC curves and AUC values obtained by AI-based identification among *BCR-ABL1* transduced/ untransduced UT-7/EPO cells with or without Mdivi-1 treatment. *BCR-ABL1*(-), *BCR-ABL1*-untransduced; *BCR-ABL1*(+), *BCR-ABL1*-transduced; Drp, dynamin-related protein; Ser, serine; Mfn, mitofusin; OPA, optic atrophy; DMSO, dimethyl sulfoxide; Mdivi-1, Mitochondrial division inhibitor 1; TEM, transmission electron microscopy; AUC, area under the curve; AI, artificial intelligence.

Given the strong possibility of Drp1 involvement in mitochondrial fragmentation in *BCR-ABL1*-transduced cells, the UT-7/EPO cells were treated with mitochondrial division inhibitor 1 (Mdivi-1), a specific inhibitor of Drp1, and mitochondrial morphology was observed using TEM. The results showed that the Mdivi-1 treatment did not alter the shape of the mitochondria in untransduced UT-7/EPO cells, suggesting that the diminished effect of Mdivi-1 in untransduced UT-7/EPO may be due to the reduced occurrence of mitochondrial fission in these cells. However, upon treatment with Mdivi-1, we observed a decrease in fragmented mitochondria instead of an increase in elongated mitochondria in *BCR-ABL1*-transduced cells (Fig. 2B).

Furthermore, we quantified the mitochondrial area through image analysis and found that Mdivi-1 treatment resulted in a slight, non-significant increase in the area per mitochondrion in *BCR-ABL1*-transduced cells (Fig. 2C). Additionally, we stained the mitochondria in these experimental groups with MitoTracker Green and observed that Mdivi-1 treatment inhibited *BCR-ABL1*-induced mitochondrial fragmentation (Fig. 2D). Furthermore, the proportion of fragmented mitochondria was significantly reduced by inhibiting Drp-1 in *BCR-ABL1*-transduced cells (p < 0.01, Fig. 2E). Moreover, the expression of p-Drp1 (Ser616) in *BCR-ABL1*-transduced cells decreased after Mdivi-1 treatment (Fig. 2F). Therefore, our findings suggest that the phosphorylation of Drp1 at Ser616 primarily mediates mitochondrial fragmentation by *BCR-ABL1*.

### AI identifies CML cells in which mitochondrial morphology is altered

After we observed that our AI technology identified *BCR-ABL1*-transduced cells and investigated mitochondrial fragmentation in these cells with different methods, further investigation was conducted to examine whether the reason for AI-based identification was the mitochondrial changes. Our results showed that two types of cells with different mitochondrial morphology, *BCR-ABL1*-transduced and untransduced UT-7/EPO cells, could be identified with high accuracy (AUC of 0.93). However, the AUC of identification between gene-untransduced cells and Mdivi-1-treated *BCR-ABL1*-transduced cells decreased to 0.78, indicating that mitochondrial morphological changes affected AI-based identification. In *BCR-ABL1*- transduced cells, the AUC was 0.81 when comparing the measurements before and after the Mdivi-1 treatment, suggesting that mitochondrial morphology between the two cells drastically changed by Mdivi-1. In contrast, the AUC of untransduced cells before and after Mdivi-1 treatment was 0.67, and the mitochondria in both cell types tended to fusion (Fig. 2G). These results suggest that AI detected morphological changes in the mitochondria in *BCR-ABL1*-transduced cells and identified *BCR-ABL1*- positive cells partly through mitochondrial alterations.

### Mitogen-activated protein kinase pathway phosphorylates Drp1 in BCR-ABL1-transduced cells

Increased Drp1 phosphorylation in *BCR-ABL1*-transduced cells suggests the activation of extracellular signal-regulated kinase (ERK) 1/2, which is reported as a responsible molecule that phosphorylates serine residues in Drp1^21^. Since ERK is also known to be activated downstream of *BCR-ABL1*, we hypothesized that ERK1/2 might mediate the p-Drp1(Ser616) observed in *BCR-ABL1*-transduced cells. Consequently, ERK1/2 phosphorylation increased in *BCR-ABL1*-transduced UT-7/EPO cells compared with that in untransduced cells (Fig. 3A). Furthermore, U0126 treatment, an inhibitor of the mitogen-activated extracellular signal-regulated kinase (MEK) 1/2 located upstream of ERK1/2, led to a concentration-dependent attenuation of ERK1/2 and Drp1 phosphorylation in UT-7/EPO cells (Fig. 3B), indicating that p-Drp1(Ser637) observed in *BCR-ABL1*-transduced UT-7/EPO is mediated through the phosphorylation of ERK1/2.

**Fig. 3.**
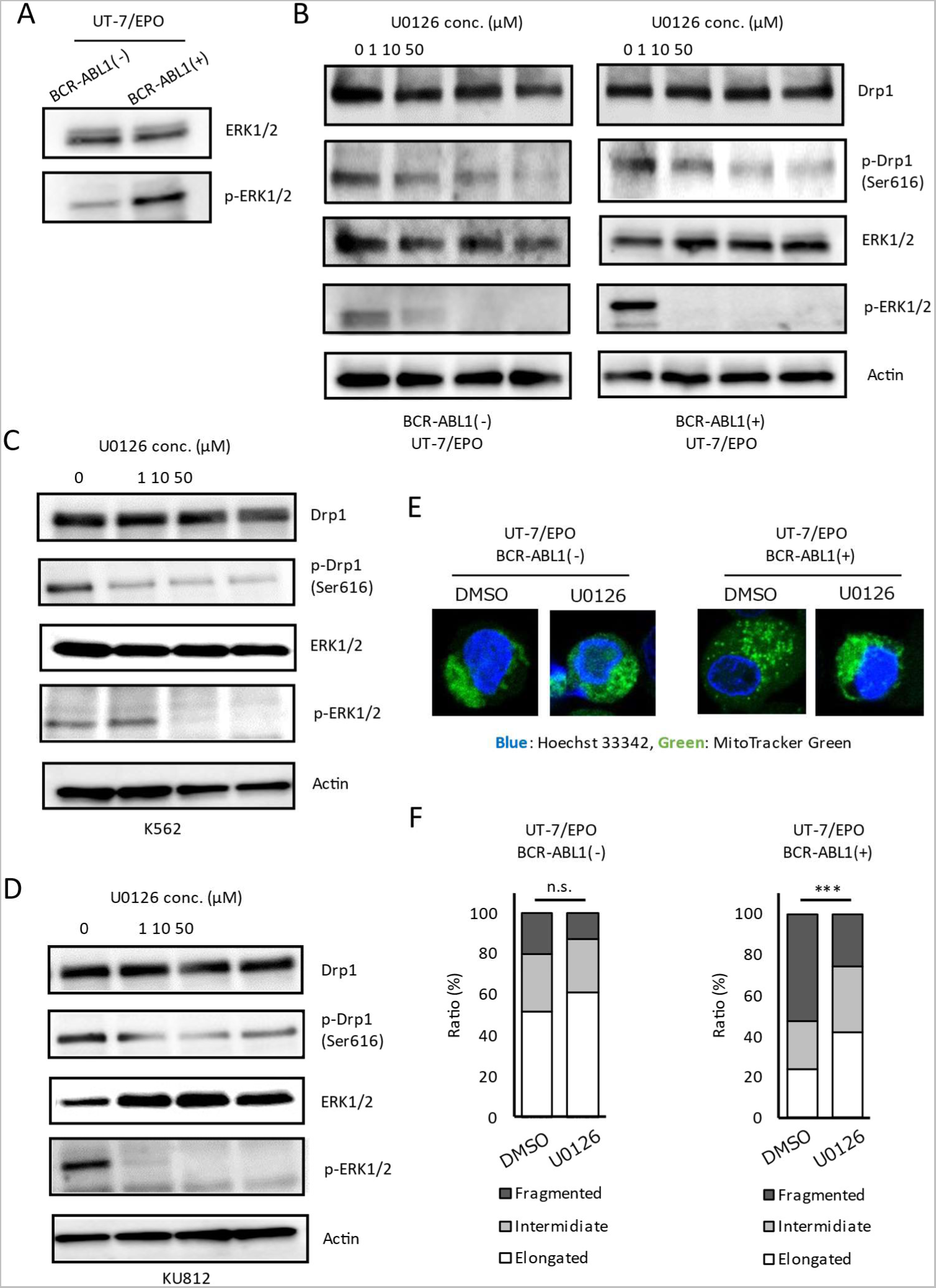
MAPK is involved in the Drp-1 phosphorylation by *BCR-ABL1*. A. ERK1/2 expression and phosphorylation of UT-7/EPO cells with or without *BCR-ABL1* transduction. B. Drp1 and ERK1/2 expression and phosphorylation of UT-7/EPO cells with or without *BCR-ABL1* transduction after treatments with 0, 1, 10, and 50 µM of the MEK inhibitor U0126 for 4 h. C. Drp1 and ERK1/2 expression and phosphorylation of K562 cells after treatments with 0, 1, 10, and 50 µM of the MEK inhibitor U0126 for 4 h. D. Drp1 and ERK1/2 expression and phosphorylation of KU812 cells after treatments with 0, 1, 10, and 50 µM of the MEK inhibitor U0126 for 4 h. E. Representative confocal laser scanning microscopy images of UT-7/EPO with or without *BCR-ABL1* treated with U0126 (10 µM for 24 h) or DMSO (control). Mitochondria are shown in green, and nuclei are shown in blue. F. The proportions of cells with different mitochondrial morphologies in UT-7/EPO with or without *BCR-ABL1* treated with U0126 (10 µM for 24 h) or DMSO (control). Among the cells imaged using confocal laser scanning microscopy, the sizes of mitochondria were classified as Elongated, Intermediate, or Fragmented, and their respective ratios were presented. ***p < 0.01 by Fisher’s exact test. *BCR-ABL1*(-), *BCR-ABL1*-untransduced; *BCR-ABL1*(+), *BCR-ABL1*-transduced; ERK, extracellular signal-regulated kinase; Drp, dynamin-related protein; Ser, serine; DMSO, dimethyl sulfoxide.

To investigate the effect of U0126 on cells that share characteristics with CML cells, we treated CML-derived K562 and KU812 cells with U0126 and observed ERK1/2 and Drp1 phosphorylation. As a result, the phosphorylation of both ERK1/2 and Drp1 was reduced in a concentration-dependent manner, similar to that in UT-7/EPO cells (Fig. 3C, D).

To observe the effect of U0126 on mitochondrial morphology, UT-7/EPO cells were treated with U0126 and observed by confocal laser microscopy. In *BCR-ABL1*-transduced cells, the fragmentation of mitochondria was suppressed by U0126 treatment (Fig. 3D), leading to a significant change in the proportion of fragmented mitochondria in response to U0126 treatment (p < 0.001, Fig. 3F).

These findings suggest that *BCR-ABL1* activates the mitogen-activated protein kinase (MAPK) pathway and enhances MAPK-mediated phosphorylation of Drp1 to induce mitochondrial fragmentation.

### CML cell proliferation was suppressed by inhibition of BCR-ABL1-induced mitochondrial fragmentation

We investigated the effect of mitochondrial fragmentation in *BCR-ABL1*-positive cells on cell proliferation. *BCR-ABL1*-transduced UT-7/EPO, K562, and KU812 cells were used as CML cell models, while gene-untransduced UT-7/EPO and acute promyelocytic leukemia (APL)-derived HL60 served as comparison targets. The effect of mitochondrial fragmentation inhibition by Mdivi-1 on cell proliferative activity was examined.

Following a 24 h stimulation with 10 µM Mdivi-1 to inhibit mitochondrial fragmentation, cell proliferative activity was significantly reduced in *BCR-ABL1*-positive UT-7/EPO, K562, and KU812 cells compared with that in the unstimulated group (p < 0.05 for all), whereas no significant difference was observed in cell proliferative activity in *BCR-ABL1*-negative UT-7/EPO and HL60 cells (p = 0.2 and 0.8, respectively, Fig. 4A). No significant difference was found in the percentage of dead cells between *BCR-ABL1*-positive and -negative cells after 24 h of stimulation (Fig. 4B).

**Fig. 4.**
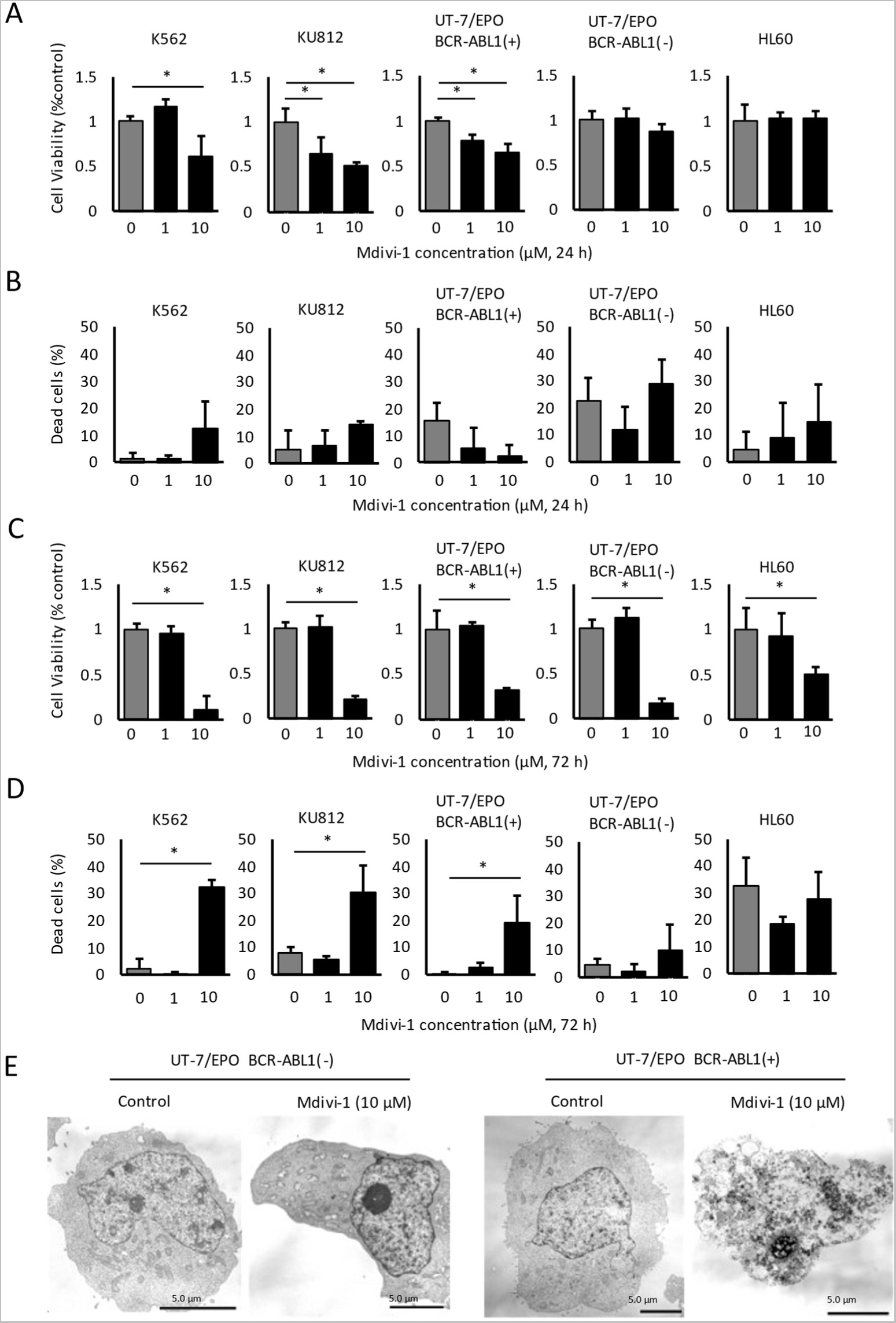
Cell viabilities and percentages of dead cells after Mdivi-1 treatment. CML-derived cell lines (K562 and KU812) and non-CML-derived cell lines (UT-7/EPO and HL60) were subjected to treatments with 1 and 10 µM of Mdivi-1 or DMSO (control) for 24 and 72 h. A. Cell viabilities were assessed after 24 h treatment of Mdivi-1 using the WST-8 assay. B. Trypan blue-positive dead cells were counted after 24 h treatment of Mdivi-1. C. Cell viabilities were assessed after 72 h treatment of Mdivi-1 using the WST-8 assay. D. Trypan blue-positive dead cells were counted after 72 h treatment of Mdivi-1. All assays were triplicated, and values are presented as the mean ± SD, *p < 0.05 by Steel’s test. E. TEM observation of *BCR-ABL1*-transduced and untransduced UT-7/EPO cell treated by 10 µM Mdivi-1. The black scale bars represent 5.0 µm. BCR-ABL1(-), *BCR-ABL1*-untransduced; BCR-ABL1(+), *BCR-ABL1*-transduced; DMSO, dimethyl sulfoxide; Mdivi-1, Mitochondrial division inhibitor 1.

Additionally, compared with that of the unstimulated group, cell proliferative activity was significantly reduced in all cell lines after 72 h stimulation with 10 µM Mdivi-1 to inhibit mitochondrial fragmentation (p < 0.05 for all, Fig. 4C). Additionally, the number of dead cells increased significantly in *BCR-ABL1*- positive UT-7/EPO, K562, and KU812 groups compared with that of the unstimulated group (p < 0.05 for all), whereas no significant increase was observed in the *BCR-ABL1*-negative UT-7/EPO and HL60 groups (p = 0.8 and 0.7, respectively, Fig. 4D). Furthermore, TEM observation revealed the induction of necrotic cell death in *BCR-ABL1*-transduced UT-7/EPO treated with Mdivi-1, which was not observed in *BCR- ABL1*-negative UT-7/EPO (Fig. 5E). These results suggest that mitochondrial fragmentation caused by *BCR-ABL1* may have a more potent effect on the proliferative activity of CML cells than *BCR-ABL1*- negative cells.

**Fig. 5.**
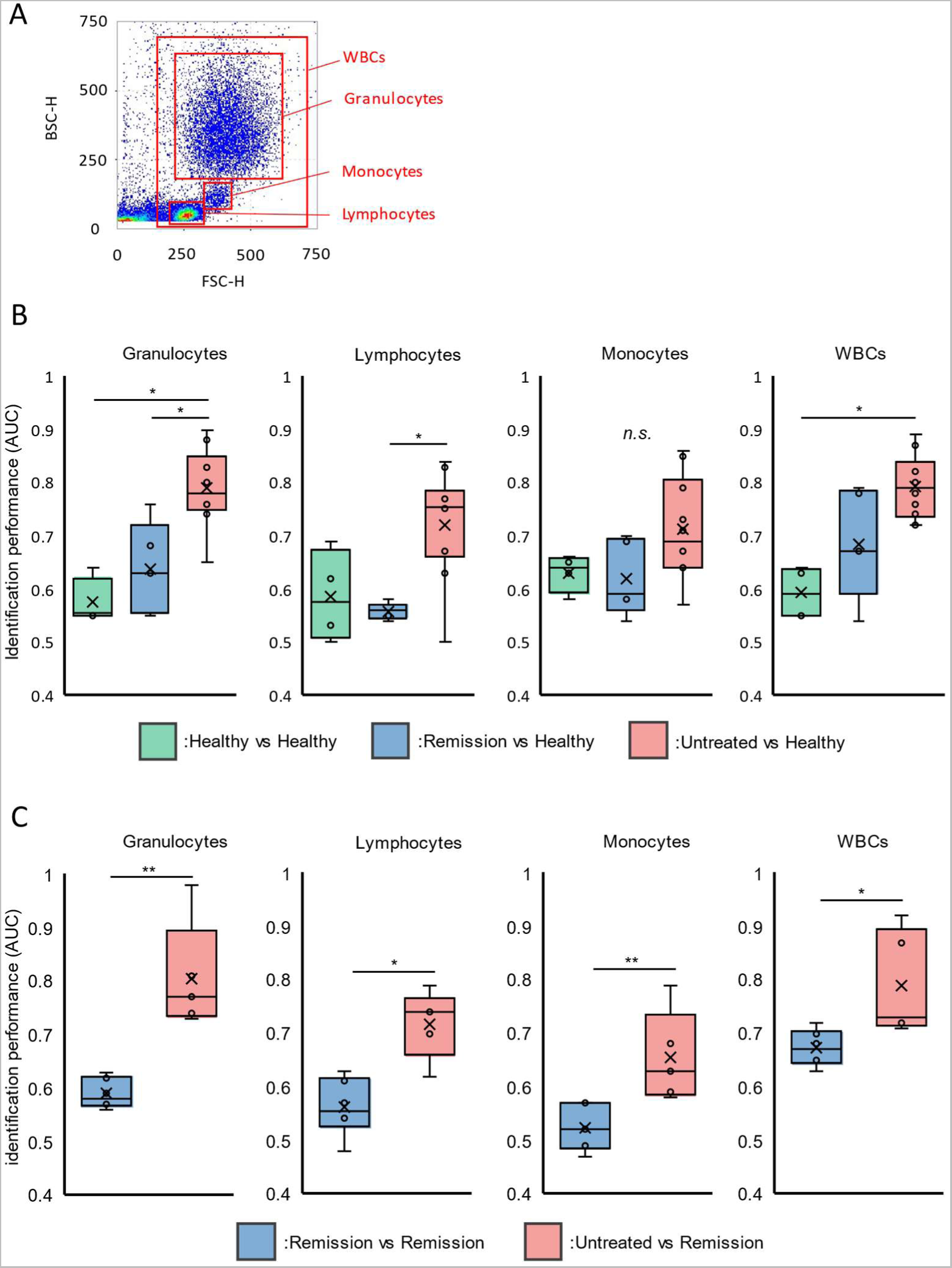
AI-based identification of peripheral white blood cells from patients with CML. A. Cells were gated into granulocytes, lymphocytes, monocytes, and WBCs from the FSC-BSC scattergrams. B. AI-based discrimination of the cells between patients with CML and healthy donors. The green, blue, and red graphs show the AUC values between two different healthy donors (n = 4), between TKI-treated remission patient with CML and a healthy donor (n = 5), and between untreated CML patients and a healthy donor (n = 10), respectively. *p < 0.05 by Steel–Dwass test; n.s., not significant. C. AI-based identifications of the cells between remission and untreated patients with CML. The blue and red graphs show the AUC values between two different remission patients (n = 6) and between remission patients and untreated patients (n = 5), respectively. *p < 0.05, **p < 0.01 by Mann–Whitney U test. AI, artificial intelligence; WBC, white blood cells; FSC, forward scatter; BSC, backscatter; AUC, area under the curve; Healthy, healthy donor; remission, CML patients who achieved remission with TKI treatment; Untreated, untreated CML patients.

### AI identifies peripheral blood leukocytes in patients with CML

After demonstrating that our AI technology could identify *BCR-ABL1*-positive cells based on features such as mitochondrial morphological changes, the next step was to evaluate the identification ability for actual leukocytes from patients with CML. Based on forward scatter (FSC) and backscatter (BSC) scattergrams, peripheral blood leukocytes were categorized into whole white blood cells (WBCs), granulocytes, lymphocytes, and monocytes (Fig. 5A). Acquired waveform signals were compared among healthy individuals (healthy), patients with CML who achieved molecular remission with TKI therapy (remission), and those with untreated CML (untreated). The results indicated that the average AUC for all cell fractions was approximately 0.6 for two different healthy individuals; the average AUC for healthy individuals and patients with CML in remission ranged from 0.6 to 0.7. In contrast, when comparing healthy individuals and untreated patients with CML, the AUC was the highest in the granulocyte and WBC fractions, 0.8 on average, which was statistically higher than that of the other comparison groups (p < 0.05 for both). In the lymphocyte fraction, the average AUC values were 0.7 (Fig. 5B).

The AUC between two different patients with CML in remission ranged from 0.5 to 0.7. In contrast, the AUC between patients with CML in remission and untreated patients with CML ranged from 0.7 to 1.0 for the granulocyte and WBC fractions (p < 0.01 and < 0.05, respectively) and from 0.6 to 0.8 for the lymphocyte and monocyte fractions (p < 0.05 and < 0.01, respectively), which were significantly higher than those of the other comparison groups (Fig. 5C). These findings suggest that AI can accurately identify peripheral blood leukocytes from untreated patients with CML and those from healthy individuals or patients with CML in remission.

### Mitochondria in peripheral blood leukocytes of patients with CML exhibit fragmentation

Since mitochondrial fragmentation was observed in cells cultured positive for *BCR-ABL1*, we examined the changes in mitochondrial morphology in leukocytes of patients with CML and found their mitochondria to be more fragmented than those of healthy individuals (Fig. 6A). In addition, imaging analysis showed a significant reduction in the area per mitochondrion in patients with CML (p < 0.05, Fig. 6B). Moreover, MMP measurements by MitoTracker DeepRed showed a decline in patients with CML compared with healthy controls in granulocytes, lymphocytes, monocytes, and WBC section as described in Fig. 5A (p < 0.05 for all), similar to the results obtained from the cultured cells (Fig. 6C, D). These results indicate that *BCR-ABL1*-induced mitochondrial fragmentation occurs in both leukocytes of patients with CML and *BCR- ABL1*-transduced cells.

**Fig. 6.**
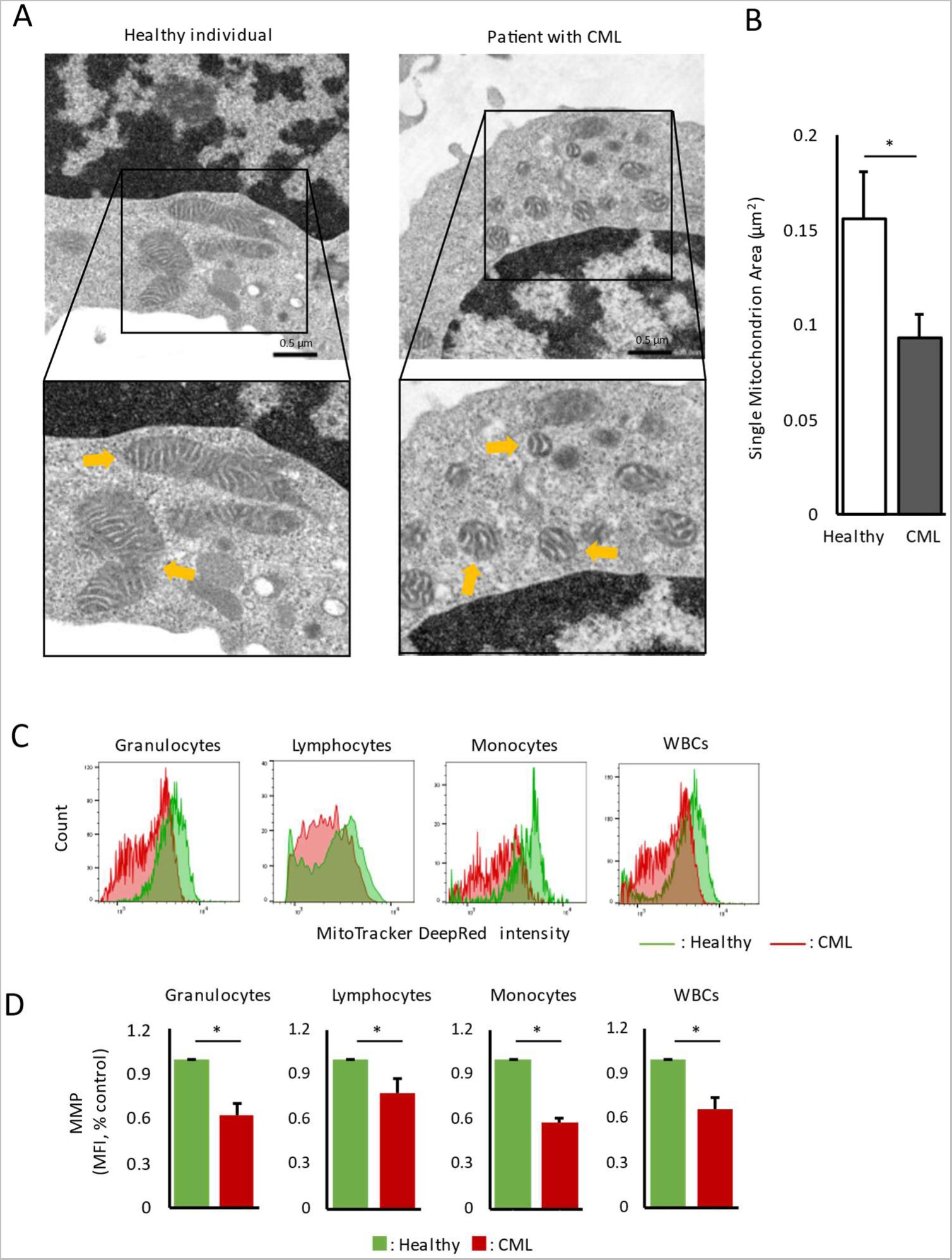
Examinations of peripheral white blood cells from patients with CML. A. Typical TEM images of peripheral white blood cells from a patient with CML and a healthy donor. The scale bars represent 0.5 µm, and the yellow arrows show the mitochondria. B. Quantitative results of single mitochondrion area of peripheral white blood cells from patients with CML (n = 4) and healthy donors (n = 3). Values are presented as the mean ± SD; *p < 0.05 by Mann–Whitney U test. C. Typical flow cytometry histogram of a healthy donor and patient with CML stained by MitoTracker DeepRed in granulocytes, lymphocytes, monocytes, and WBC fraction. D. MMP measurement of peripheral white blood cells from patients with CML (n = 3) and healthy donors (n = 3) from each cell fraction. The average MFI of MitoTracker Deep Red staining was compared. Values are shown relative to those of the healthy donors. Values are presented as the mean ± SD; *p < 0.05 by Mann–Whitney U test. TEM, transmission electron microscopy; WBC, white blood cells; MMP, mitochondrial membrane potential; MFI, median fluorescent intensity; Healthy, healthy donor; CML, untreated CML patients.

## Discussion

This study investigated the feasibility of peripheral CML cell detection and identified morphological changes in CML cells as a potential basis for AI-mediated identification, with the aim of early-stage diagnosis of CML by detecting small amounts of CML cells using our novel AI technology. Our findings suggest that AI could effectively identify *BCR-ABL1*-positive cells without any labeling in both model cell lines and real-world clinical specimens. Additionally, our data demonstrated that *BCR-ABL1* amplified mitochondrial fragmentation by augmenting the MAPK-mediated phosphorylation of Drp1, which we considered a potential basis of the AI-based identification of CML cells (Extended data 1).

The modulation in GC-derived waveform signals reflects structural changes within the cells^22^. Since the dot diameter inferred from the optical arrangement of the structured illumination was 0.7 µm^18^, in theory, GC can detect structures larger than 0.7 µm. Mitochondria, essential intracellular organelles, can range several micrometers in size and are found in large quantities throughout the cytoplasm^23^. Mitochondria have a high refractive index directly proportional to the membrane potential^24^, indicating that they strongly influence the waveform signals. Previous reports^18, 19, 22^ on GC have not sufficiently discussed the basis for its identification; strikingly, our comprehensive study strongly suggests that our AI technology could detect changes in organelles such as mitochondria and allowed us to uncover a portion of the “black box”^25^. Moreover, the AI may have captured structural changes other than mitochondrial alterations in CML cells; therefore, further studies regarding this aspect are warranted.

Mitochondria are widely recognized for their role as energy-producing organelles that generate adenosine triphosphate (ATP); however, they are also involved in numerous other crucial cellular functions, such as apoptosis induction, metabolism regulation, reactive oxygen species production, redox homeostasis maintenance, and calcium ion regulation^26^. In recent years, mitochondrial dynamics, which involve morphological changes caused by mitochondrial fission and fusion, have garnered significant attention^27^. Mfn1/2 are involved in mitochondrial outer-membrane fusion, whereas OPA1 mediates inner membrane fusion^28^. Mitochondrial fission is stimulated in the cytoplasm by Drp1, which localizes to the outer mitochondrial membrane via phosphorylation at Ser616 while inhibiting fission via phosphorylation at Ser637^29^. Changes in mitochondrial dynamics have been implicated in various cellular functions and disease pathogeneses. For example, effector T-cell mitochondria tend to undergo fission, whereas memory T-cell mitochondria tend to undergo fusion^30^. In many solid tumors, including lung, colon, and breast cancers, mitochondrial fragmentation has been linked to metabolic reprogramming, cell cycle progression, and increased migration, invasion, and metastatic potential^31^. Interestingly, it has been reported that mitochondrial fragmentation is involved in the pathogenesis of myelodysplastic syndromes, and that mitochondrial morphogenesis may represent a potential therapeutic target^32^. Our findings indicate that the inhibition of *BCR-ABL1*-induced mitochondrial fragmentation suppresses cell proliferation, suggesting that the altered mitochondrial dynamics observed in CML cells may be involved in CML cell function and pathogenesis. These results suggest that the inhibition of Drp1 expression represents a novel potential therapeutic strategy for CML.

To the best of our knowledge, this is the first report revealing morphological and functional alterations in CML cells. Additionally, MAPKs are involved in mitochondrial fragmentation and may play some role in CML pathogenesis. As reported previously, c-Abl enhances Drp1 phosphorylation and mitochondrial fragmentation in neurons^33^. Thus, in CML cells, mitochondrial fragmentation may be enhanced by the constant Abl activity induced by the BCR-ABL1 fusion protein. Reports on ABL1 activation of MAPK in CML cells^34^ and MAPK-induced Drp1 phosphorylation in Huntington’s disease cells^21^ are consistent with our results. In practice, MAPK inhibition has been reported to suppress the proliferation of CML cells^35^ and enhance sensitivity to TKIs^36^, while MEK inhibition has been shown to be effective in blast-phase CML treatment^37^. Thus, alterations in mitochondrial function mediated by MAPK in CML have been suggested as potential therapeutic targets. The low rate of TFR achievement has been attributed to the persistence of residual CML-leukemia stem cells (CML-LSCs), even after TKI treatment^10, 38^. New treatment targets for CML-LSCs may directly contribute to increasing the rate of TFR achievement. While many studies have aimed at eradicating CML-LSCs^39–41^, few have aimed at early diagnosis and treatment initiation to improve the TFR achievement rate.

Another intriguing finding of this study was the accurate identification of unlabeled CML cells by AI in both cell line models and actual patient cells. In general, CML is diagnosed based on increased WBC counts. In daily clinical settings, more than 90% of neutrophils are already positive for *BCR-ABL1* based on fluorescent *in situ* hybridization. However, existing automated blood cell analyzers and clinical examinations cannot detect CML cells in the early stage of the disease, and the WBC count is generally within the normal range. Our results suggest that AI-based identification, along with routine laboratory tests, such as complete blood counts, may enable the detection of minute quantities of CML cells in peripheral blood during screening. This approach could potentially enhance the TFR success rate by facilitating early diagnosis and treatment initiation of CML.

A limitation of this study is that the identification performance of AI between CML and other hematologic disorders was not verified. Thus, it remains unclear whether AI-based CML cell identification is specific to CML or not. Additionally, this study only tested the discriminative performance between healthy and CML specimens where almost all cells were *BCR-ABL1*-positive, and further validation of the analysis algorithm is necessary to detect small amounts of CML cells.

In summary, we showed that the AI-based high throughput technology was employed to detect cells exhibiting the morphological changes in the mitochondria implicated in CML pathogenesis, and label-free identification of CML cells in peripheral blood can be achieved using the technology.

## Methods

### Cell culture

K562, KU812, and HL-60 cells were cultured in RPMI 1640 medium (Nacalai Tesque, Tokyo, Japan) supplemented with 10% heat-inactivated fetal bovine serum (FBS; Nichirei, Tokyo, Japan) and penicillin-streptomycin (100 U/mL; Nacalai Tesque), in a humidified incubator maintained at 37°C under 5% CO_2_ conditions. The UT-7/EPO cells^42^ were cultured in Iscove’s modified Dulbecco’s medium (Nacalai Tesque) supplemented with 10% heat-inactivated FBS, 100 U/mL penicillin-streptomycin, and 10 ng/mL recombinant human erythropoietin (EPO, Peprotech, Cranbury, NJ, USA), in a humidified incubator maintained at 37°C under 5% CO_2_. To achieve *BCR-ABL1* expression in UT-7/EPO, cDNAs were subcloned into the pMSCV-IRES-green fluorescence protein vector (Addgene #20672) with reference to a report by Masubuchi et al.^43^ The constructed plasmid was validated by sequencing prior to use.

### Clinical samples

Blood samples were obtained from 10 patients with CML without any prior treatment, 17 patients with CML who had achieved molecular remission through TKI treatment, and 5 healthy volunteers. This study was approved by the ethics committee of Juntendo University School of Medicine (M12-0895-M01) and Sysmex Corporation (2021-068) and conducted in adherence with the Declaration of Helsinki. The patients provided informed consent by accessing the study information published on the website and were free to withdraw their consent at any point during the study, as per the Ethical Guidelines for Medical and Health Research Involving Human Subjects. Therefore, the ethics committee waived the requirement for written informed consent. The clinical information and data pertaining to clinical tests were retrieved from the medical records (Extended data 2).

### AI analysis using GC

To collect whole blood samples, red blood cells were lysed in five times the amount of red blood cell lysis solution (Qiagen, Hilden, Germany), followed by 15 min of incubation at room temperature (24°C). Subsequently, white blood cells were washed in a phosphate-buffered solution (PBS) (Nacalai Tesque) containing 2% FBS. Then, 1.0 × 10^6^ cells were incubated at room temperature for 20 min with 5 µL of APC-labeled anti-CD45 antibody (Clone: 2D1, BioLegend, San Diego, CA, USA). The cell lines were washed in PBS containing 2% FBS, followed by staining with MitoTracker Deep Red (10 nM; Thermo Fisher Scientific, Waltham, MA, USA) for 30 min at 37°C in a humidified incubator.

AI analysis was performed using a prototype product based on GC technology developed by ThinkCyte K.K., Tokyo, Japan. The protocol was performed according to the manufacturer’s instructions. Based on forward scatter (FSC) and backscatter (BSC) scattergrams, BSC-low/FSC-low cells were defined as lymphocytes, BSC-mid/FSC-mid cells as monocytes, and BSC-high/FSC-high cells as granulocytes. One of these cells was stained as described above and analyzed using machine learning to compare the identification performance between the two groups of cells. Fluorescence waveform signals from 2,000 cells were utilized to train a model based on a support vector machine (SVM) algorithm. Furthermore, waveform signals from other 500 cells were utilized to verify the trained model. The waveforms from each cell were scored for classification based on the SVM model. (Extended data 3). The true positive and false positive rates of the classification results for each cell were used to create receiver operating characteristic (ROC) curves. The identification results were assessed using the area under the curve (AUC) of the ROC curves.

### Wright–Giemsa staining

The cells were immobilized onto glass slides using Cytospin (Sakura Finetek, Tokyo, Japan), followed by staining with Wright’s solution (Muto pure chemicals, Tokyo, Japan) containing 50% methanol (Nacalai Tesque) for 5 min. Subsequently, the cells were washed and stained with Giemsa solution (Muto pure chemicals) for 20 min. The cells were then washed, dried, and observed under a microscope (ECLIPSE TE300, Nikon, Tokyo, Japan).

### Transmission electron microscopy

For TEM, the cells were fixed with 2.5% glutaraldehyde (TAAB, Berkshire, UK) in 0.1 M phosphate buffer (pH 7.4), followed by post-fixation with 2% OsO_4_ in the same buffer. The fixed specimens were dehydrated using a graded ethanol series and embedded in Epok812 (Oken shoji, Tokyo, Japan). Ultrathin sections were prepared and stained with uranyl acetate (Wako, Tokyo, Japan) and lead citrate (TAAB). These sections were examined using a transmission electron microscope (JEM-1400 Flash, JEOL, Tokyo, Japan). For the quantification of mitochondrial areas using the ImageJ software^44^ (version 1.53t), mitochondria on the TEM images were encircled, and the selected areas were quantified.

### Confocal laser scanning microscopy

One million cells were incubated with MitoTracker Green (100 nM; Thermo Fisher Scientific) for 30 min in a humidified incubator maintained at 37°C, followed by Hoechst 33342 staining (1 µg/mL; Dojindo, Kumamoto, Japan) for 5 min at room temperature. Next, cells were washed, immobilized onto glass slides, and observed using LSM 780 (Zeiss, Oberkochen, Germany) with a water-immersion objective lens at 64-fold magnification. Short and rounded mitochondria were defined as fragmented, and cells with fragmented mitochondria accounting for 70% or more of the total mitochondrial population were classified as ’Fragmented,’ while cells with fragmented mitochondria accounting for 30% or less were classified as ’Elongated.’ Cells with an intermediate ratio were classified as ’Intermediate.’ A total of 100 cells were counted in each group, and the proportions were compared using Fisher’s exact test.

### MMP

One million cells were incubated with MitoTracker Deep Red (10 nM; Thermo Fisher Scientific) for 30 min in a humidified incubator maintained at 37°C. Following a wash with PBS, the mean fluorescent intensity (MFI) was quantified using flow cytometry (FACSCelesta, Becton Dickinson, Franklin Lakes, NJ, USA). Data were analyzed using FlowJo software (version 10.8.1, Becton Dickinson).

### Western blotting

The cells were exposed to either Drp1 inhibitor Mdivi-1 (1, 10, and 50 µM for 24 h; Tokyo Chemical Industry, Tokyo, Japan) or MEK inhibitor U0126 (1, 10, and 50 µM for 4 h; Promega, Madison, WI, USA). A corresponding volume of dimethyl sulfoxide (Nacalai Tesque) was added as a control.

To obtain total proteins, 5.0 × 10^5^ cells were lysed in radioimmunoprecipitation assay buffer (Nacalai Tesque) for 30 min on ice. The samples were then subjected to sodium dodecyl sulfate polyacrylamide gel electrophoresis, transferred onto a polyvinylidene difluoride membrane (Millipore, Bedford, MA, USA), probed with antibodies, detected with Western BLoT Rapid Detect v2.0 (Takara, Shiga, Japan), and developed using Western BLoT Chemiluminescent HRP Substrate (Takara). The chemiluminescent images were observed using FUSION FX (Vilber, Paris, France) and FUSION software (version 17.03, Vilber). The following antibodies were employed: anti-DRP1 Ab (Clone: D8H5, Cell Signaling Technology, Danvers, MA); anti-phospho Drp1 (Ser616) Ab (Cell Signaling Technology); anti-Mitofsin1 Ab (Clone: 11E91H12, Abcam, Cambridge, UK); anti-Mitofsin2 Ab (Clone: D2D10, Cell Signaling Technology); anti-OPA1 Ab (Clone: 18/OPA-1, Becton Dickinson); anti-ERK1/2 antibody (clone W15133B, BioLegend); anti-ERK1/2 phospho(Thr202/Tyr204) antibody (clone: 6B8B69, BioLegend); and anti-Actin Ab (clone: 13E5, Cell Signaling Technology).

### Cell viability assay

An appropriate medium was used to culture 5.0 × 10^3^ cells, with or without Mdivi-1, at concentrations of 1 and 10 µM (Tokyo Chemical Industry). Following incubation for 24–72 h, cell viability was assessed by measuring the 450 nm absorbance of reduced WST-8 (2-(2-methoxy-4-nitrophenyl)-3-(4-nitrophenyl)-5-(2,4-disulfonyl)-2H-tetrazolium, monosodium salt) using the Cell Counting Kit-8 (Dojindo). Moreover, dead cells were counted using TC200 (BioRad, Benicia, CA, USA) after staining the cells with trypan blue (Wako).

### Statistical analyses

AI-based identification results between the two types of cells were evaluated using AUC values. AUC was represented on a scale between 0.5 and 1.0; the value increased as the identification results improved. For statistical analyses, the Mann–Whitney U test was used to compare differences between two experimental groups, and the Steel–Dwass test was used to compare differences among more than three experimental groups. Steel’s test was used to compare differences between the control and other groups. Fisher’s exact test was used to compare differences in proportion between two experimental groups. All statistical tests were two-tailed, and p < 0.05 indicated statistical significance. Statistical analyses were performed using EZR, which is based on R and R commander^45^. In boxplots, the values were represented by the 10th, 25th, 50th (median), 75th, and 90th percentiles, and individual values were superimposed on the boxplots using black symbols.

## Acknowledgements

This work was funded by Sysmex Corp. and ThinkCyte K.K.. to T.T. as a collaborative research project. We thank Kaho Shioiri from the Department of Hematology, Juntendo University Graduate School of Medicine, for providing us with the *BCR-ABL1*-transduced UT-7/EPO cell lines. We also thank the Laboratory of Morphology and Image Analysis, Cell Biology, Proteomics and Biomolecular Science, and Molecular and Biochemical Research in Biomedical Research Core Facilities, Juntendo University Graduate School of Medicine, for their technical assistance. We would like to sincerely thank Dr. Fumimaro Takaku from Jichi medical school for his considerable advice and ideas for this study.

## Author contributions

K.S. designed, performed, and analyzed experiments and contributed to writing the manuscript. N.W. collected clinical specimens and information and contributed to clinical discussions. S.To. contributed to scientific discussions. S.A. analyzed TEM images and contributed to scientific discussions. K.O. contributed to clinical and technical discussions. S.Ts. collected clinical specimens and information. K.Y. supervised the project. Y.K. and S.O. provided methodology. N.K., S.S., and M.A. supervised the research and contributed to clinical and scientific discussions. T.T. designed and supervised the research, collected clinical specimens and information, and contributed to writing the manuscript. All authors approved the final manuscript.

## Data availability statements

The datasets generated and/or analyzed during the current study are available from the corresponding author upon reasonable request.

