## Extended data for "Artificial Intelligence Enables the Label-Free Identification of Chronic Myeloid Leukemia Cells with Mitochondrial Morphological Alterations"

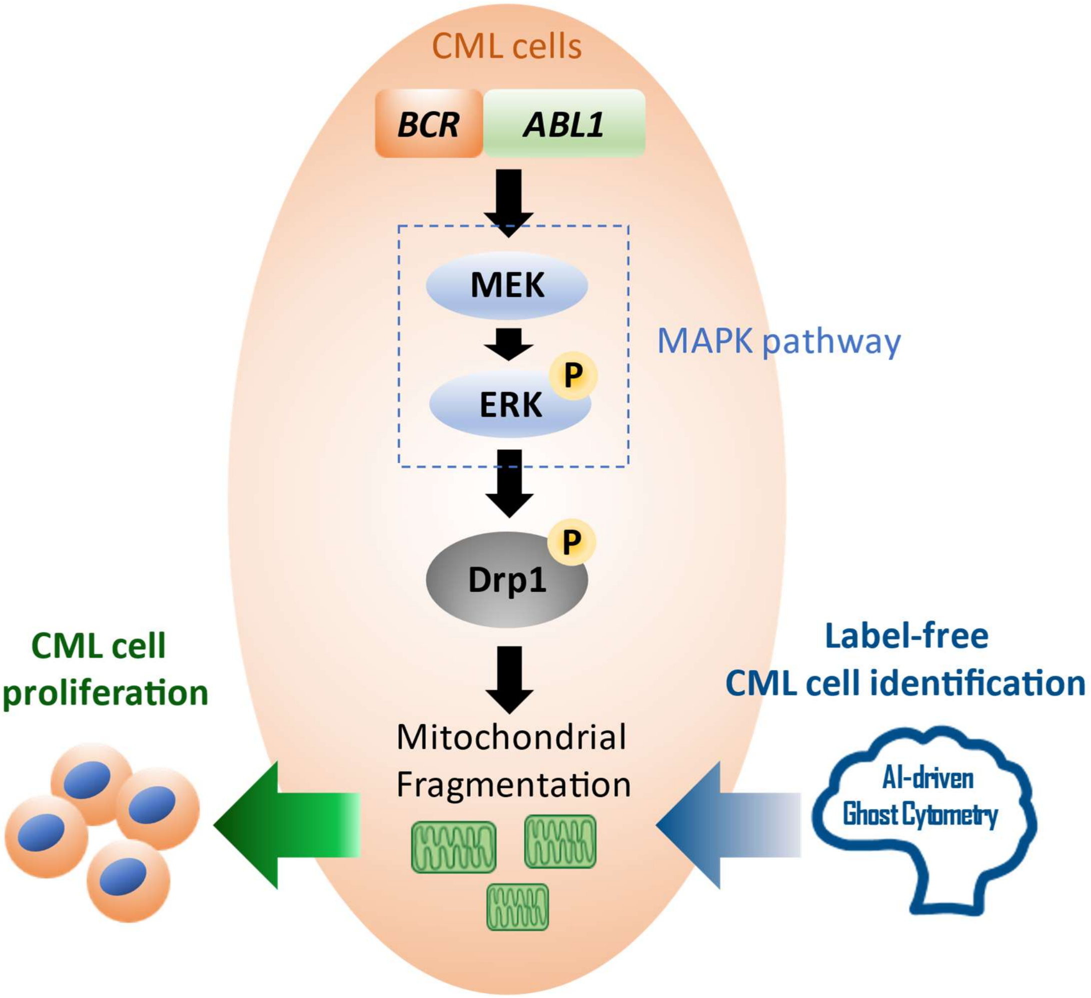


Extended data 1:

An overview of the study is presented. BCR-ABL1 induces mitochondrial fragmentation by upregulating the phosphorylation of Drp1 through the MAPK pathway. AI can identify BCR-ABL1-positive CML cells in a label-free manner by detecting the morphological changes including those in mitochondria. Mitochondrial fragmentation was considered to involve in CML cell proliferation.

CML, chronic myeloid leukemia; Drp, dynamin-related protein; MEK, mitogen-activated extracellular signal-regulated kinase; ERK, extracellular signal-regulated kinase; AI, artificial intelligence.

**Extended data 2.** Patient characteristics

| | Untreated patients with CML (n = 10) Median (IQR) | Patients with CML in remission (n = 17) Median (IQR) | P-value |
|---|---|---|---|
| Age (years) | 63.5 (53.5–68) | 65 (48–73) | P = 0.73 |
| Sex (M/F) | 7/3 | 11/6 | N/A |
| WBC ($10^9$/L) | 86.7 (54.3–133.4) | 6 (4.7–6.9) | P < 0.001 |
| RBC ($10^{12}$/L) | 3.8 (3.6–4.4) | 4.3 (4.1–4.5) | P = 0.21 |
| PLT ($10^9$/L) | 439 (352–617) | 247 (168–265) | P < 0.05 |
| N-band (%) | 28.6 (19.7–37.1) | 3 (2–5) | P < 0.001 |
| N-seg (%) | 38 (30.2–44.1) | 67 (50–76.5) | P < 0.001 |
| Lympho (%) | 3.5 (2.7–8.6) | 24 (16.5–30.5) | P < 0.001 |
| Mono (%) | 2.4 (1.5–5.2) | 4.5 (3–6) | P = 0.57 |
| Eosi (%) | 3.4 (1.1–5.1) | 2 (2–3) | P = 0.20 |
| Baso (%) | 4.2 (2.4–4.8) | 0 (0–0.5) | P < 0.001 |
| Blast (%) | 0 (0–0) | 0 (0–0) | N/A |
| Promyelo (%) | 0.4 (0–1.2) | 0 (0–0) | P < 0.05 |
| Myelo (%) | 11.6 (7.3–18.1) | 0 (0–0) | P < 0.001 |
| Meta (%) | 0.6 (0.2–1.7) | 0 (0–0) | P < 0.001 |
| BCR-ABL IS% | 102.1 (93.3–111) | 0.0023 (0–0.02) | P < 0.001 |

Characteristics of patients with CML without any TKI treatments (n = 10) and those who achieved deep molecular remission (n = 17). Values are shown in median (interquartile range; IQR). P-values were calculated by Mann–Whitney U test.

CML, chronic myeloid leukemia; TKI, tyrosine kinase inhibitor; WBC, white blood cells; RBC, red blood cells; PLT, platelet; N-band, band neutrophils; N-seg, segmented neutrophils; Lympho, lymphocytes; Mono, monocytes; Eosi, eosinophils; Baso, basophils; Promyelo, promyelocytes; Myelo, myelocytes; Meta, metamyelocyte; BCR-ABL IS%; international scale of *BCR-ABL1* mRNA level; N/A, not applicable.

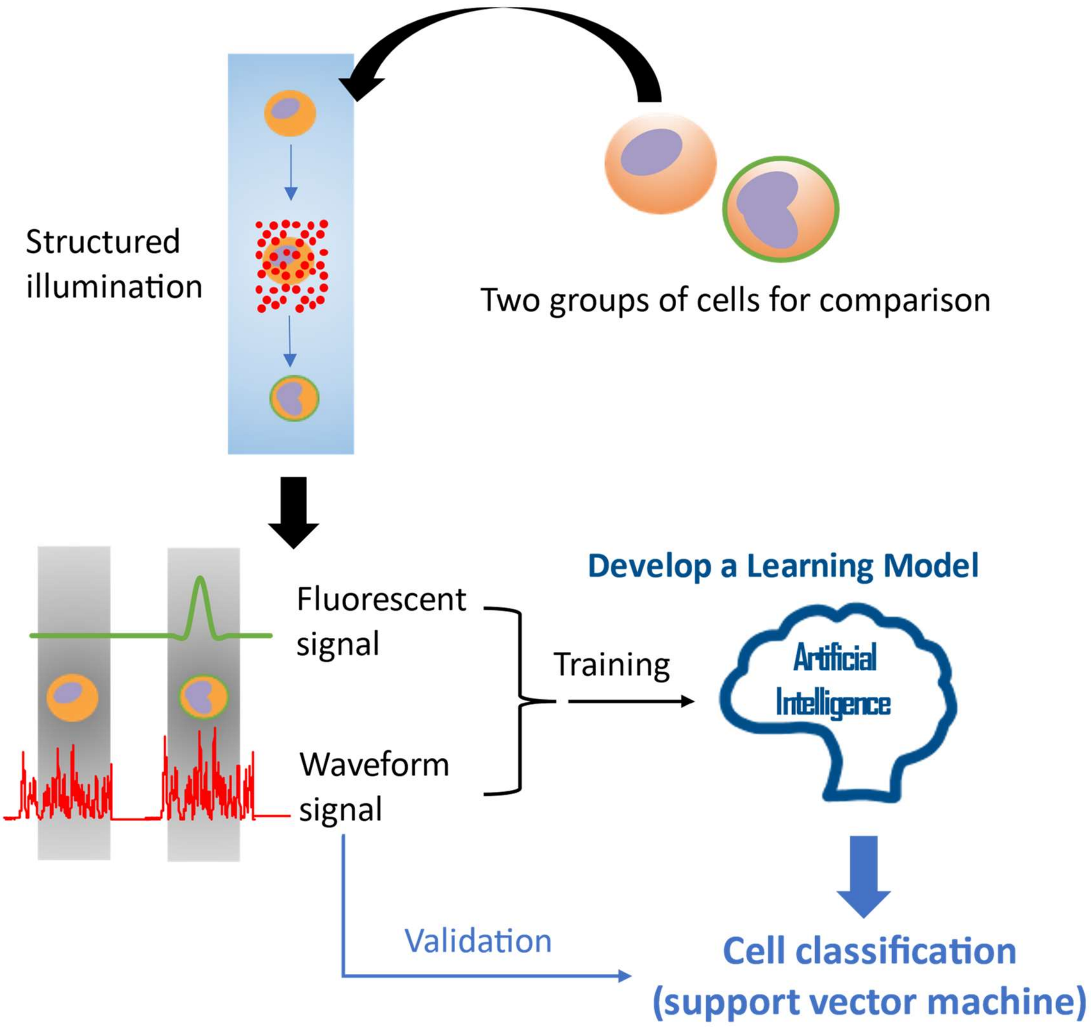


Extended data 3:

A methodology outline for the ghost cytometry (GC) analysis is presented. One of the two cell groups selected for comparison was labeled, combined, and analyzed using a machine learning-integrated high-content flow cytometry. Structured illumination provides multiparametric temporal waveform signals that reflect subtle structural changes within the cells. Fluorescence and label-free waveform signals obtained from a sample of 2000 cells were employed to develop a machine learning model using a support vector machine method. Subsequently, only the waveform signals obtained from 500 additional cells were used to validate the trained model.
